# Distinct Control States Underlie Voluntary Task Switching: Evidence for Capacity-Dependent Control Modes

**DOI:** 10.64898/2026.04.12.716959

**Authors:** Hyung-Bum Park, Monica Rosenberg, Edward Vogel

## Abstract

People often switch tasks when attention wanes or an alternative task becomes more appealing. Such choices may reflect different control modes that may vary with working memory (WM) capacity. This study tested whether momentary attentional lapses prospectively predict voluntary task switching and whether this relationship depends on WM capacity. Participants performed a continuous performance task involving face and scene images, with blocks in which they either freely chose the next task or followed an externally imposed sequence. The results suggested a capacity-dependent pattern in which individuals with lower WM capacity were more likely to switch following lapse-prone blocks, whereas higher-capacity individuals tended to switch from relatively well-focused states. Eye-tracking revealed greater bias toward the competing irrelevant category before switches in lower-capacity individuals, accompanied by early conflict-related pupil dilation. Externally imposed task sequencing selectively reduced lapses in the lower-capacity group without affecting higher-capacity performance, suggesting that external structure can scaffold weaker internal goal maintenance. These findings suggest that the relationship between lapses and voluntary switching varies with WM capacity rather than being uniform across individuals. This pattern is consistent with a goal-competition account in which lapses reflect shifts in the balance between competing task goals, and voluntary switches may be preceded by different control states.

---

The ability to sustain attention over extended periods is essential for successful goal-directed behavior, yet even simple tasks produce frequent lapses in control. In laboratory settings, these transient failures are indexed by commission errors in continuous performance tasks (CPT), which serve as sensitive markers of momentary failures in executive regulation (Robertson et al., 1997). Performance on sustained attention tasks typically declines over time on task, and this decline is accompanied by dynamic changes in large-scale neural control networks (Langner & Eickhoff, 2013; Esterman et al., 2014; Massar et al., 2016).

Traditional explanations for attentional lapses have converged on two major theoretical accounts. Resource depletion or overload accounts propose that sustained performance taxes limited attentional resources. This increases variability and lapses as control demands accumulate and is a view that is well established in the vigilance literature (Kahneman, 1973). In contrast, underload accounts emphasize that low task demands or automatized performance fail to engage attention, allowing mind wandering and task-irrelevant thoughts to intrude (McVay & Kane, 2012; Thomson et al., 2015). Although these perspectives differ in mechanism, both conceptualize lapses as passive control failures. However, motivational and reward manipulations often leave the time on task decline relatively intact and lapses also occur in engaging contexts, suggesting limitations for purely depletion or underload perspectives (Unsworth & McMillan, 2013).

A growing body of work offers a complementary theoretical perspective and reframes attentional lapses as the consequence of competition among simultaneously active goal representations. According to this *goal-competition* framework, off-task states emerge when alternative goals gain sufficient strength to challenge and disrupt the current task set, thus diverting cognitive resources (Smallwood & Schooler, 2015). Recent theoretical work links sustained attention to task switching by conceptualizing prolonged focus as the maintenance of one task set in the presence of competing alternatives. This parallels accounts of mind wandering that emphasize concurrent goal conflict (Oberauer, 2024). Within this perspective, lapses reflect transient dominance of alternative mental sets rather than solely weakened control.

Task-switching paradigms provide a natural experimental framework to examine the goal-competition hypothesis because they instantiate competition between task sets. Classic findings show reliable switch costs, manifested as slower responses and increased errors after a switch. These costs are typically interpreted as reflecting the cognitive effort required to reconfigure the active task set and shield it from interference by the previously relevant task set (Monsell, 2003; Kiesel et al., 2010). These retrospective analyses of switch costs, however, do not reveal whether lapse-related control states preceding voluntary switches carry predictive signatures of emerging goal conflict. Recent work using a switch-based CPT provides an important bridge between sustained attention and task switching. Chidharom and colleagues (2025) used a go/no-go CPT with bilateral face and scene stimuli in which participants were cued every 20 trials to repeat or switch task goals. Instructed switches were treated as periods of heightened goal competition as the previously relevant task set could interfere with the newly relevant one. They found increased commission errors early after instructed switches relative to repeat blocks, suggesting that CPT lapses can reflect competition between task goals rather than nonspecific vigilance failure. Building on this finding, the present study takes a complementary prospective approach by asking whether lapse-related markers of goal competition predict subsequent voluntary decisions to abandon the current task set.

Voluntary task-switching paradigms are well suited to address this question because individuals autonomously choose which task to perform next. Such choices reveal intrinsic preferences shaping endogenous goal selection and highlight how people balance persistence against exploration (Arrington & Logan, 2004). Emerging evidence suggests that voluntary switching reflects cost-benefit evaluations related to exploration-exploitation trade-offs in adaptive control (Fröber et al., 2019; Fröber & Dreisbach, 2016; Wilson et al., 2014). Yet the precise control mechanisms that give rise to voluntary switches remain unclear. If lapses are signatures of competition between concurrently active goals, a natural prediction would be that periods of heightened goal-competition should manifest as increased lapse rates and should prospectively predict subsequent choices to abandon the current task in favor of an alternative.

The goal-competition perspective aligns with individual difference approaches to cognitive control. The dual mechanisms of control framework proposes that people vary in their reliance on proactive control, which sustains goal representations in anticipation of conflict, or on reactive control, which engages after interference is detected (Braver, 2012). Critically, individuals vary systematically in their reliance on these modes, with working memory (WM) capacity serving as a key determinant of control strategy preference (Kane & Engle, 2003).

High-capacity individuals better maintain goal representations and resist distraction, whereas lower-capacity individuals rely more on reactive adjustments and are more vulnerable to goal neglect and task-inappropriate responses (Engle, 2010). Short-timescale links between sustained attention fluctuations and WM maintenance suggest shared control substrates (deBettencourt, Keene, Awh, & Vogel, 2019). Moreover, individual differences in sustained attention also covary with arousal regulation, with lower capacity associated with dysregulated arousal that contributes to lapses (Unsworth & Robison, 2017). Large-scale functional connectivity profiles predict trait-like variation in sustained attention, underscoring stable differences in goal maintenance (Rosenberg et al., 2016). Accounts of meta-control describe how cognitive systems balance stability, which protects the current goal against interference, and flexibility, which enables updating when demands change, and they situate WM capacity as a factor that shifts this balance (Goschke & Bolte, 2014; Fröber & Dreisbach, 2021). Importantly, meta-control can operate proactively, preparing appropriate control states based on contextual cues, and reactively, adjusting control after detecting the need for change (Kang & Chiu, 2021).

The present task extended this logic by embedding the CPT within a task-switching context in which competing goals were present on every trial. In a modified spatial go/no-go CPT, participants viewed one face and one scene stimulus presented bilaterally, but only one category was task-relevant within a given block. Thus, each display contained both the currently relevant task category and a competing, irrelevant category. This design allowed us to measure two lapse-related error types that captured different levels of control failure. Commission errors indexed failures to withhold a response on no-go trials, providing a response-level marker of transient control failure. Swap errors indexed responses directed to the currently irrelevant category, providing a more task-specific marker of goal confusion because they indicated that behavior was guided by the competing task set rather than the currently relevant one. In the free-choice session, participants chose which task to perform next after each block, allowing these markers of goal competition to be measured prospectively before voluntary task choices were made. In addition, eye tracking captured covert competition through the proportion of initial saccades directed to the irrelevant category, and phasic pupil dilation served as a physiological readout of cognitive effort and control engagement (van der Wel & van Steenbergen, 2018; Aston-Jones & Cohen, 2005).

By contrasting blocks that preceded voluntary switches with those that preceded repetitions, the present study adopts a prospective approach to cognitive control, examining whether momentary fluctuations in lapse-related performance predict subsequent task selection. Within a goal-competition framework, lapses are not treated as passive failures but as indicators of shifts in balance between competing task representations. Thus, the central prediction was that lapse-related markers of goal competition in the current block would prospectively relate to participants’ subsequent voluntary task choices. We further examined the theoretically motivated possibility that this prospective relation may vary as a function of WM capacity. One possibility is that individuals with lower WM capacity may be more likely to switch reactively following lapse-prone states that reflect reduced goal maintenance, whereas individuals with higher capacity may switch from relatively well-focused states, consistent with stronger goal shielding and more deliberate exploration.

To further assess the role of control demands in task selection, a forced-choice session was included in which task sequences were externally imposed. This manipulation allows us to test whether decision autonomy itself imposes additional cognitive demands, motivated by decision fatigue that arises when faced with excessive choices (e.g., Netflix paradox; see Iyengar & Lepper, 2000). Prior work shows that imposing external structure can mitigate cognitive demand and support control, particularly for individuals with limited resources (i.e., lower WM capacity) by outsourcing the cognitive demand associated with maintaining and evaluating competing goal representations (Risko & Gilbert, 2016). Therefore, this provides a critical test of whether the observed switching patterns reflect differences in control mode rather than differences in overall performance level.

In summary, our approach represents a fundamental shift from a retrospective to a prospective analysis of cognitive control dynamics. Rather than inferring control states from post-switch performance costs, we examine the antecedent signatures that predict voluntary decisions while they are still forming. This forward-looking perspective captures the temporal unfolding of goal competition and reveals how momentary fluctuations in attentional control shape higher-level decisions about task engagement (Smallwood & Schooler, 2015).

## Method

### Participants

Twenty-nine volunteers (13 female, 16 male) participated in the experiments and received monetary compensation ($20 per hour). Participants were aged between 18 and 31 years (*M* = 23.7, *SD* = 3.6), reported normal or corrected-to-normal visual acuity, and provided informed consent according to procedures approved by the University of Chicago Institutional Review Board. Sample size was determined via an a priori power analysis (G*Power 3.1; Faul et al., 2009), targeting 85% power to detect a medium effect size (Cohen’s *d* > 0.5) in primary within-subject comparisons between task switching conditions (i.e., pre-switch vs. pre-stay blocks) at α = .05, based on prior studies investigating attentional lapses in task-switching CPT (Chidharom et al., 2025). This power analysis was designed for the primary within-subject contrast between task conditions and was not designed to estimate power for the cross-level interactions in which between-subject differences in WM capacity moderate within-subject effects.

### Stimuli and Procedure

Stimuli were generated in MATLAB (The MathWorks, Natick, MA, USA) using the Psychophysics Toolbox (Brainard, 1997) and displayed on an LCD computer screen (BenQ XL2430T; 120 Hz refresh rate; 24.5 inch diagonal; 1,920 × 1,080 pixels) with a uniform grey background (15.1 cd/m²) and positioned approximately 70 cm from participants. Eye movements were recorded with a desk-mounted EyeLink 1000 Plus (SR Research) sampling at 1000 Hz. For analysis, gaze and pupil time series were down-sampled to 100 Hz. The experiment comprised two parts: (1) a WM change-detection task to measure individual WM capacity, and (2) a modified spatial go/no-go CPT with inter-block task selection. Figure 1 illustrates the stimuli and overall task procedure.

**Figure 1.**
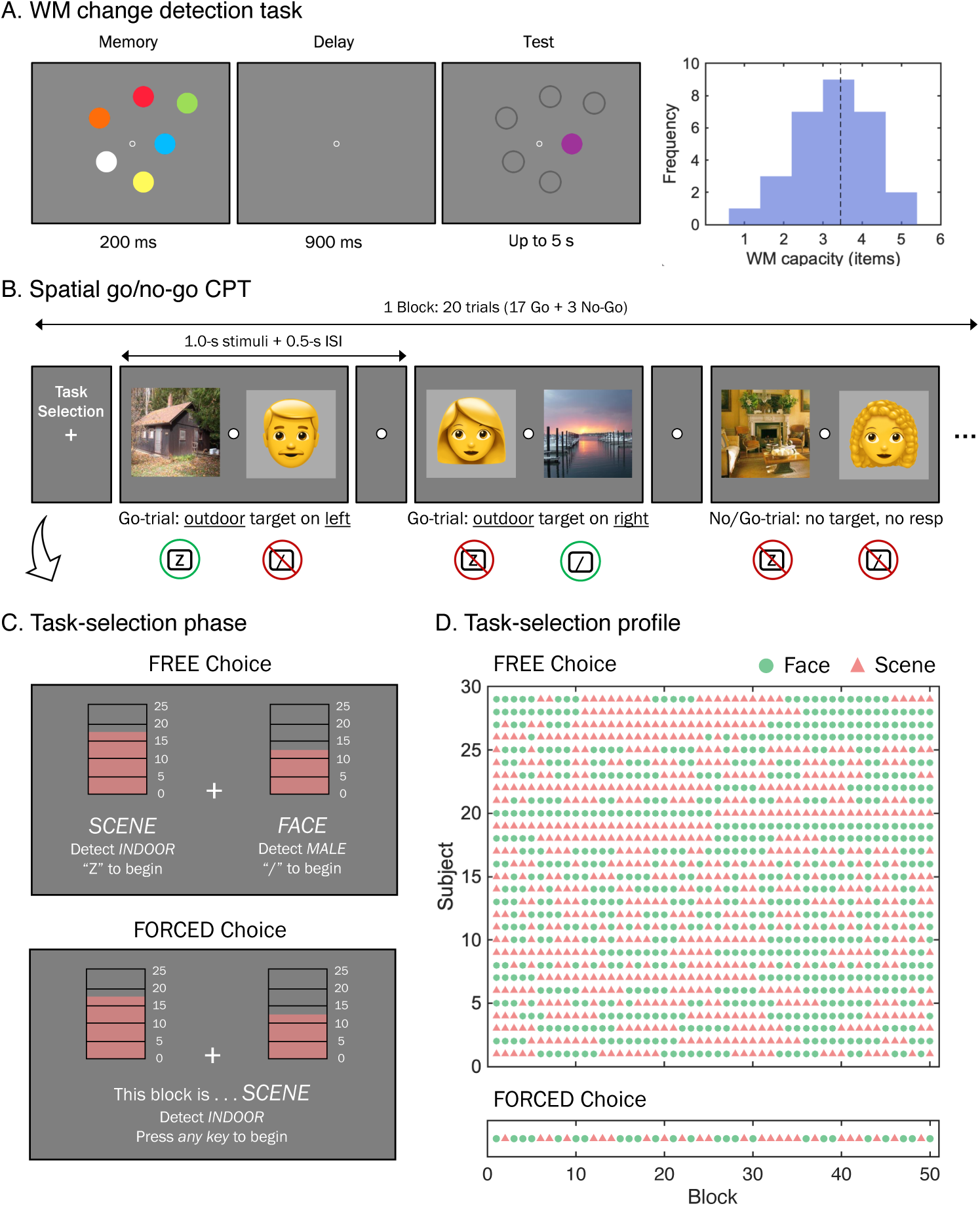
Experimental design and task overview. (A) Working memory (WM) change-detection task used to estimate individual capacity (Cowan’s K). Right panel shows distribution of WM capacity across participants. (B) Modified spatial go/no-go continuous performance task. Example shows the scene task with outdoor targets. Each trial presented one face and one scene stimulus. Participants reported the block-defined target location on go trials (85%) and withheld responses on no-go trials (15%). Each block contained 20 trials and began with a task-selection phase. (C) Inter-block task-selection display, showing block gauges with remaining blocks for each task type. In the free-choice session (top), participants autonomously selected next task. In the forced-choice session (bottom), computer program assigned next task after 3 s countdown. (D) Task-selection patterns across 50 blocks, showing subject-by-subject task sequences in the free-choice session (each row is participant), and a fixed pseudo-random sequence used in the forced-choice session.

#### Working Memory Change-Detection Task

Prior to the main CPT sessions, all participants completed a color change-detection task to assess individual WM capacity (Figure 1A). Each trial began with a central fixation cross (0.4° visual angle) presented for 500 ms, followed by a memory array of six colored squares (2.0° × 2.0° each) presented for 200 ms. The memory items appeared at randomly selected locations within an imaginary circle of 10.0° radius, with a minimum center-to-center distance of 2.7° to prevent crowding effects. After a 900 ms retention interval, a test array appeared in which one randomly selected item either remained the same color or changed to a new color (50% probability each). Participants indicated whether any color had changed by pressing the “z” key for no-change or the “/” key for change.

Participants completed 80 trials total (40 change, 40 no-change). Colors were selected from eight categorically distinct hues with high discriminability (red, green, blue, magenta, yellow, cyan, orange, and white). Cowan’s *K* was calculated using the standard formula (Cowan, 2001): K = (hit rate – false alarm rate) × set size. This measure estimates the number of items maintained in WM, with higher K values indicating greater capacity.

#### Modified Go/No-Go Continuous Performance Task

Following the WM assessment, participants completed two sessions of a modified CPT incorporating both face and scene stimuli (Figure 1B). Each session comprised 50 blocks (25 scene task blocks, 25 face task blocks) with task type determined by the experimental manipulation described below. Scene stimuli were sampled from the SUN image database (Xiao et al., 2010), including 995 indoor scenes (e.g., kitchens, bedrooms, offices, etc.) and 1,341 outdoor scenes (e.g., landscapes, streets, parks, etc.). Face stimuli were sampled from the Chicago Face Database (Ma et al., 2015), selecting 363 female and 351 male faces of diverse ethnicities displaying neutral expressions. All images were randomly presented to participants.

The modified CPT employed a spatial go/no-go paradigm in which participants made location discrimination responses to target stimuli while withholding responses when targets were absent. Each trial presented one face and one scene image (5.7° × 5.7°) simultaneously at 7.0° eccentricity to the left and right of central fixation for 1000 ms, with left/right positioning counterbalanced across trials. A central fixation cross (0.4°) remained visible throughout stimulus presentation. Following stimulus offset, a 500 ms blank interval preceded the next trial. Participants indicated the spatial location of target stimuli using keyboard responses (“z” for left position, “/” for right position) but only when targets were present. Speeded responses were encouraged but responses were allowed from stimulus onset until the end of the subsequent blank interval (1500 ms total response window).

Task rules were defined at the block level. For face task blocks, participants responded to a predetermined target gender, either male or female faces. For scene task blocks, participants responded to a predetermined scene type, either indoor or outdoor scenes. Target assignments (male vs. female faces; indoor vs. outdoor scenes) were counterbalanced across participants and remained constant throughout the experiment for each individual. Go trials, in which the designated target was present, occurred on 85% of trials (17 of the 20 trials within a block). The remaining 15% were no-go trials, in which the relevant category was present but did not match the target exemplar. The correct response on these trials was to withhold any keypress. A *commission* error was defined as a response made on a no-go trial and was treated as the primary index of attentional lapses. Because each display always included both a relevant and an irrelevant category stimulus, erroneous responses directed toward the irrelevant item were separately coded as *swap* errors, which serve as a direct behavioral marker of goal confusion and indicate misallocation of attention between competing task sets. Each block consisted of 20 trials, lasting approximately 30 s.

At the end of each block, participants entered a task-selection phase that differed by session (Figure 1C). In the *free-choice* session, participants chose whether to perform the next block as a face task or a scene task. They were shown a “block gauge” displaying the number of remaining blocks for each task type, along with a reminder of the target category. Pressing the “z” key initiated the scene task, while pressing “/” initiated the face task. Participants were informed that they needed to complete all 25 blocks of each task, but they could freely determine the order across the 50 blocks in this session. Once all blocks of one task type were completed, that option was no longer available.

In the *forced-choice* session, participants were informed which task they would perform next according to a predetermined pseudorandom sequence rather than making their own choice. The same block gauge and target reminders appeared on screen, followed by a 3-s countdown period. The computer then revealed the assigned next task. Participants pressed either key to proceed to the designated block. The forced sequence was identical for all participants and contained 30 switches distributed across the 50 blocks. Session order (free-choice vs. forced-choice) was counterbalanced across participants. All tasks were completed in a single laboratory visit. The forced-choice and free-choice sessions each lasted approximately 35 minutes, and the full study took approximately 90 minutes to complete, including eye-tracker calibration, the WM change-detection task, practice trials, and both CPT sessions.

### Data Analysis

All behavioral and physiological analyses included WM capacity as a key moderating variable, with subjects assigned to high or low capacity groups according to a median split of their Cowan’s *K* estimates. Post-hoc analyses confirmed a substantial difference in capacity estimates between the two groups, with high-*K* subjects (*N* = 14, *M* = 4.11 items, *SD* = 0.47) and the low-*K* subjects (*N* = 15, *M* = 2.55 items, *SD* = 0.75).

#### Behavioral Responses

Behavioral indices were computed per block. Commission error rate was the proportion of responses on no-go trials. Swap error rate was the proportion of responses to the irrelevant category. For the free-choice session, each block was labeled by the choice taken immediately afterward (*n*−1 task selection), resulting in blocks followed by a voluntary switch labeled “pre-switch”, and blocks followed by a voluntary repetition labeled “pre-stay”. Accordingly, the blocks after the choice (*n*+1) were labeled “post-switch” or “post-stay”, respectively. Task selection was unconstrained at the beginning of the 50 block sequence but gradually became more constrained as the completed blocks approached the total of 25 blocks per task. By design, the final block of each session is therefore determined by the remaining task type. Note that the same block can serve as a post-choice block for decision *n* and as a pre-choice block for decision *n*+1. For the forced-choice session, the same post-choice (*n*+1) comparisons were computed, but because order was externally imposed, analyses focused on blocks following imposed switch versus repetition.

We modeled the probability of a voluntary switch on block n+1 from behavioral performance on block *n* using binomial generalized linear mixed-effects model (GLME). Trials were first aggregated to the block level within participant. For each block, commission error rate was computed as the proportion of responses on no-go trials, and swap error rate was computed as the proportion of responses directed to the currently irrelevant category. We then constructed within-subject lagged predictors so that switch probability of block *n*+1 was predicted from commission and swap error rates on block *n*. The first block for each participant was excluded from this analysis because there was no preceding block from which lagged performance predictors could be computed.

The analysis dataset therefore comprised, for each participant and block, a binary outcome indicating whether the next block was a voluntary switch (*Switch*_*n*+1_∈{0, 1}) and two lagged error predictors defined on the preceding block, commission proportion (*Commission_n_*) and swap proportion (*Swap_n_*), together with WM capacity (*K*). All continuous predictors were z-scored to standardize scale. The GLME model took the form:

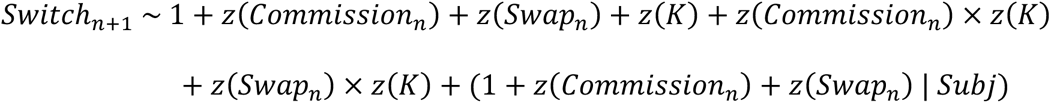

In this model, random effects were estimated at the participant level, including a random intercept and random slopes for the two within-subject lapse predictors. This allowed participants to differ in their overall tendency to switch as well as in the extent to which prior-block commission and swap errors predicted subsequent switch probability. The fixed interaction terms tested whether these within-participant lapse–switch associations varied systematically with between-subject differences in WM capacity. Models were fit by maximum likelihood with a logit link using the Laplace approximation. We report fixed-effect estimates on log-odds with 95% confidence intervals (CI_95%_) and odds ratios where informative.

#### Eye-Tracking Data

Eye-tracking analyses were conducted within the same framework, with the goal of testing whether misallocation of attention to task-irrelevant stimuli during a block predicted the subsequent decision to switch tasks in the free-choice session. Eye-position samples were drawn from trial-level gaze recordings (100 Hz) and expressed relative to the central fixation point (0, 0). Gaze samples were then blink-corrected (velocity spike filter; samples during blinks ±50 ms removed) and linearly interpolated.

Two complementary measures captured distinct aspects of visual goal selection. First, we quantified an initial-saccade index by computing, for each block, the proportion of trials in which the first saccade during stimulus presentation landed on a distractor belonging to the currently irrelevant category. Second, to examine the overall spatial distribution of gaze, we analyzed fixation positions along the horizontal axis. To standardize across bilateral displays, trials were spatially mirrored so that the target stimulus always appeared on the left side and the distractor on the right. We then estimated kernel density functions separately within each block, generating subject-level difference curves contrasting pre-switch and pre-stay blocks. These curves were averaged within groups stratified by working memory capacity using a median split on Cowan’s *K*. To assess the reliability of between-group differences in gaze distribution, we applied a subject-level bootstrap with 5,000 resamples combined with spatial clustering. Contiguous regions along the horizontal dimension where the CI_95%_ of the bootstrap difference excluded zero were considered statistically reliable.

Pupil traces were converted to relative change within trial by dividing by the mean of the first five post-onset samples and subtracting one, then time-locked to stimulus onset following established protocols for cognitive pupillometry (Mathôt et al., 2018). For the prospective contrast, switch-minus-stay difference traces were formed from the preceding block per subject and averaged within WM groups for display. Statistical inference used cluster-based permutation tests with 5,000 iterations on unsmoothed traces from 0 to 1,500 ms, with samples taken every 10 ms. This approach allowed identification of temporally contiguous periods during which pupil dilation significantly differentiated switch versus stay conditions, accounting for multiple comparisons.

### Transparency and Openness

We report how sample size was determined, all manipulations, all measures, and data exclusions (if any) in the study. The study design and analyses were not preregistered. All data generated or analyzed during this study are available via the Open Science Framework repository at https://osf.io/j3457/. ChatGPT (OpenAI) was used to assist with language editing and proofreading, and identification of potentially relevant literature during manuscript preparation. All scholarly claims, analyses, citations, and references were reviewed and confirmed by the authors.

## Results

### Behavioral Overview of Voluntary Switching

Across the 50 blocks in the free-choice session, participants made voluntary task switches on average 13.2 times, CI_95%_ [9.5, 16.8], shown in Figure 1D. Switching frequencies did not differ reliably between high-*K* (*M* = 14.6, CI_95%_ [10.7, 18.6]) and low-*K* participants (*M* = 11.8, CI_95%_ [5.7, 17.9]), *t*(27) = 0.76, *p* = .456, Cohen’s *d* = 0.14, and also did not differ by session order, with comparable switch counts for participants who completed the free-choice session first (*M* = 13.1, CI_95%_ [8.4, 17.8]) and those who completed the forced-choice session first (*M* = 13.3, CI_95%_ [6.5, 20.1]), *t*(27) = −0.06, *p* = .954, Cohen’s *d* = −0.01. Thus, overall switching tendency was comparable across WM capacity groups and session-order groups. The critical question, however, is *when* participants chose to switch. As detailed below, prospective analyses show that the internal control state that precedes a voluntary switch is shaped by WM in a manner consistent with a goal-competition account and the dual-mechanisms distinction between proactive and reactive control.

### Prospective Markers of Voluntary Switching

Figure 2 summarizes the behavioral results. We asked whether lapse-related CPT indices in block *n* prospectively differentiated the choice for block *n+*1, and whether this relation depended on individual’s WM capacity. Lapses were indexed by commission errors (“go” | no-go) and swap errors (responses to irrelevant category). We first examined this question using a two-way mixed-effects analysis of variance (ANOVA) on behavioral indices of lapses as a function of pre-choice block (pre-switch vs. pre-stay) and WM capacity group (high-*K* vs. low-*K*).

**Figure 2.**
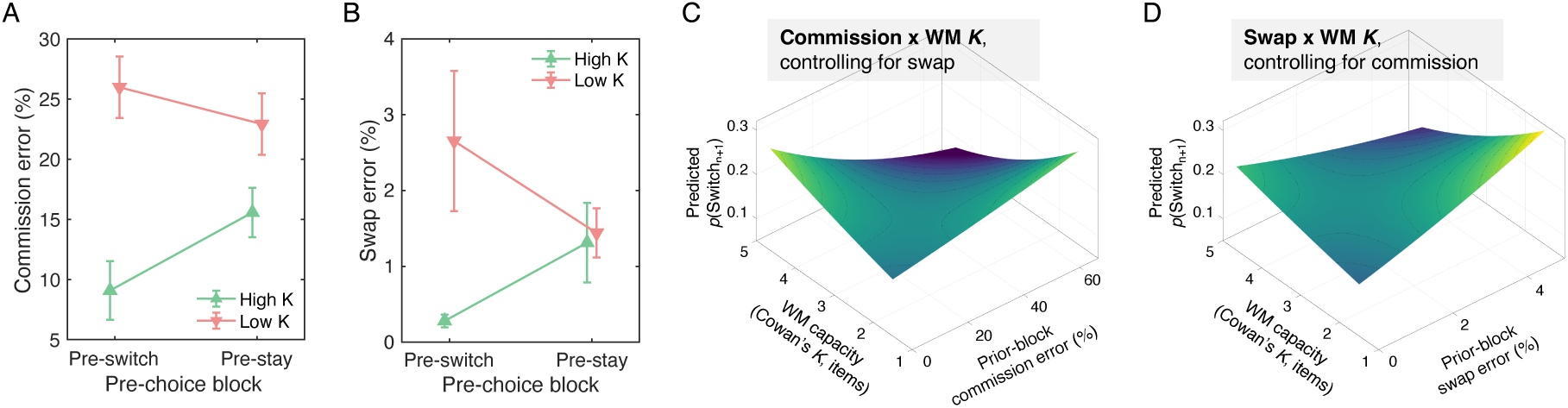
Behavioral markers of lapses and goal confusion prospectively predict voluntary task switching in a capacity-dependent manner. (A) Commission error rates (responses on no-go trials) in blocks immediately preceding voluntary task choices, plotted by upcoming decision (pre-switch vs. pre-stay) and working memory (WM) capacity group using a median split on Cowan’s *K*. (B) Swap error rates (responses to currently irrelevant category), plotted using the same format. Error bars represent standard error of the mean. (C) Predicted switch probability from the simultaneous binomial generalized linear mixed-effects model showing the unique *Commission × K* interaction effect after controlling for swap error rate. (D) Predicted switch probability from the same model showing the unique *Swap × K* interaction effect after controlling for commission error rate. The model predicted switch likelihood on block *n+*1 from lagged commission and swap error rates on block *n*, continuous WM capacity estimates, and their interactions, with subject-specific random slopes for both within-subject error predictors.

For commission errors (Figure 2A), the crossover pattern was reflected in a significant two-way interaction between pre-choice block and WM capacity group, *F*(1, 27) = 5.72, *p* = .024, η_p_² = .18. Low-K participants committed 26.0%, CI_95%_ [21.0, 31.0], commission errors in pre-switch blocks versus 22.9%, CI_95%_ [17.9, 27.9], in pre-stay blocks, *t*(14) = 1.06, *p* = .305, Cohen’s *d* = 0.28. High-K participants showed the opposite pattern, with 9.1%, CI_95%_ [4.3, 14.1], commission errors in pre-switch blocks versus 15.6%, CI_95%_ [11.5, 20.6], in pre-stay blocks, *t*(13) = −2.36, *p* = .035, Cohen’s *d* = −0.65. The main effect of pre-choice block was not significant, *F*(1, 27) = 0.74, *p* = .399, η_p_² = .03, whereas the main effect of WM capacity group was significant, *F*(1, 27) = 18.96, *p* < .001, η^p^² = .41, reflecting overall higher commission error rates in the low-K group.

Swap errors (Figure 2B), as a behavioral marker of task-level goal confusion in our modified spatial go/no-go CPT, also revealed a significant block-by-group interaction, *F*(1, 27) = 5.19, *p* = .031, η_p_² = .16. Low-*K* participants showed 2.7%, CI_95%_ [0.8, 4.5] swap errors before switches versus 1.4%, CI_95%_ [0.8, 2.1] before stays, *t*(14) = 1.50, *p* = .156, *d* = 0.40, whereas high-*K* participants showed 0.28%, CI_95%_ [0.1, 2.1] before switches versus 1.31%, CI_95%_ [0.3, 2.0] before stays, *t*(13) = −1.92, *p* = .077, *d* = −0.53. The main effect of pre-choice block was not significant, *F*(1, 27) = 0.03, *p* = .856, η_p_² = .00, but the main effect of WM capacity group was marginal, *F*(1, 27) = 3.87, *p* = .059, η_p_² = .13, with overall higher swap errors in low-K group.

To treat Cowan’s *K* estimates as a continuous moderator, we fit a binomial GLME predicting the probability of a voluntary switch on block *n*+1 from the lagged commission and swap error rates on block *n* and WM capacity. Commission and swap errors were entered simultaneously as predictors, with subject-specific random slopes included for both error measures. The simultaneous inclusion of both commission and swap error rates was motivated by an initial examination of their overlap. At the trial level, only 5.4% of commission errors were also coded as swap errors, and about 11.5% of swap errors were also coded as commission errors. Thus, most commission errors reflected responses on no-go trials that were not directed to the irrelevant task category, whereas most swap errors reflected task-category confusion that did not occur on no-go trials. At the block level, prior-block commission and swap error rates were positively but modestly correlated, *r* = .24, CI_95%_ [.19, .29], *p* < .001. These results indicate that the two measures were related but not redundant, supporting their inclusion as simultaneous predictors of voluntary switch probability. The analysis included 1,421 block transitions.

At the mean of *K* and controlling for swap errors, higher prior-block commission error rate predicted lower odds of switching on the subsequent task-selection, *b* = −0.24, CI_95%_ [−0.45, −0.04], *p* = .019. The main effect of swap error rate was not significant at the mean of *K*, *b* = −0.162, CI_95%_ [−0.40, 0.08], *p* = .188, nor was the main effect of *K*, b = 0.01, CI_95%_ [−0.46, 0.47], *p* = .979. Critically, both error measures interacted reliably with WM capacity. The *Commission × K* interaction remained significant when controlling for swap errors, *b* = −0.23, CI_95%_ [−0.43, −0.03], *p* = .022. The *Swap × K* interaction was also significant when controlling for commission errors, *b* = −0.64, CI_95%_ [−1.06, −0.22], *p* = .003. As shown in Figure 2C and D, both interactions reflected a capacity-dependent reversal in the relation between prior-block errors and subsequent switching. That is, commission and swap errors were associated with relatively higher switch probability at lower *K* but lower switch probability at higher *K*.

Together, these behavioral results indicate that voluntary switches may be preceded by qualitatively different internal states as a function of WM capacity. Low-*K* participants tended to switch from lapse-prone states consistent with reactive control, whereas high-*K* participants tended to switch from well-focused states consistent with proactive goal-shielding.

### Oculomotor Signatures of Goal Competition

Gaze provided converging evidence for the same capacity-dependent goal selection. The proportion of trials in which the initial saccade landed on the currently irrelevant-category distractor showed the crossover observed in the error indices (Figure 3A). A two-way mixed ANOVA with pre-choice block and WM capacity group again revealed a significant interaction, *F*(1, 27) = 9.92, *p* = .004, η_p_² = .27. Descriptively, low-*K* individuals directed their first saccade to the distractor on 4.5%, CI_95%_ [3.1, 6.0] of trials in pre-switch blocks versus 3.3%, CI_95%_ [2.0, 4.5] in pre-stay blocks, *t*(14) = 1.82, *p* = .091, Cohen’s *d* = 0.49. High-*K* individuals showed the opposite tendency with 1.9%, CI_95%_ [0.5, 3.4] in pre-switch blocks versus 3.4%, CI_95%_ [1.8, 4.7] in pre-stay blocks, *t*(13) = −2.82, *p* = .015, Cohen’s *d* = −0.78. The main effect of pre-choice block and WM capacity group were not significant, *Fs*(1, 27) < 1.69, *ps* > .205, η_p_²s < .06.

**Figure 3.**
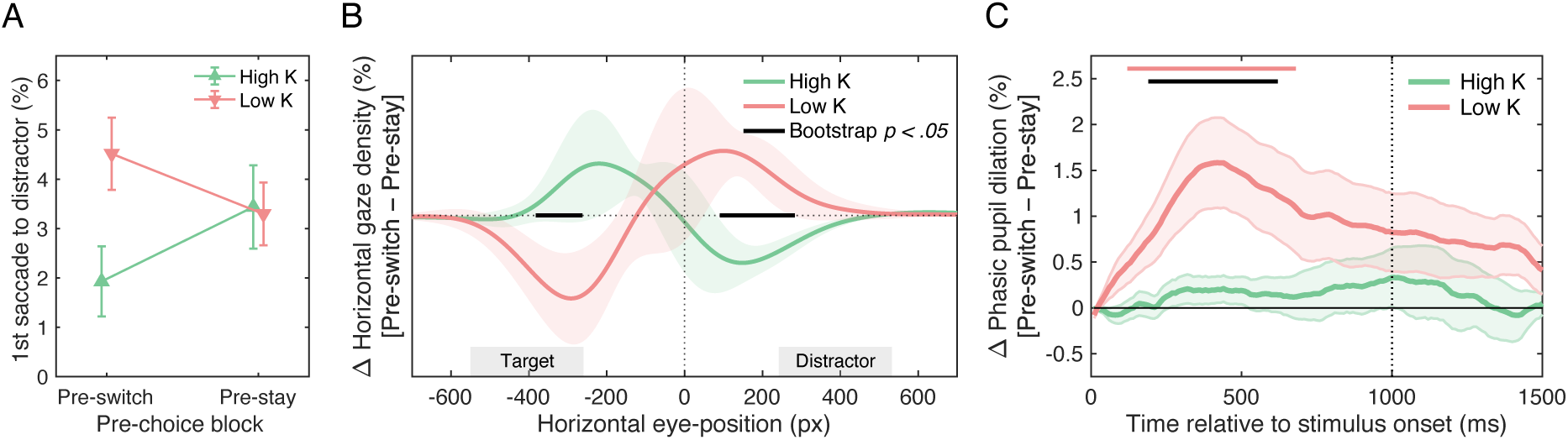
Oculomotor and pupillary markers reveal capacity-dependent goal competition. (A) Initial saccade bias toward irrelevant category distractor, showing crossover interaction by upcoming decision (pre-switch vs. pre-stay) and working memory capacity group (High *K* vs. Low *K*). Error bars represent standard error of the mean. (B) Horizontal eye-position density differences (pre-switch minus pre-stay) across bilateral stimulus display, plotted separately for low-capacity (red) and high-capacity (green) groups. Shaded areas represent 95% confidence intervals. Black bars indicate ranges where bootstrap analysis revealed reliable between-group differences (*p* < .05). (C) Stimulus-locked phasic pupil dilation difference traces (pre-switch minus pre-stay), normalized within participant. Red and black bar on top indicate significant within-group and between-group contrasts, respectively.

A fixation-density analysis further contrasted horizontal eye-position distributions between pre-switch and pre-stay blocks (Figure 3B). Group-mean difference curves showed a double-dissociation pattern that mirrors the behavioral interactions. Subject-level bootstrap with temporal clustering identified reliable between-group divergence over a *target*-defined region (−262 to −382 px) and a *distractor*-defined region (90 to 283 px) where the CIs_95%_ excluded zero. High-*K* group showed greater relative density near the target and reduced density near the distractor before switching, whereas low-*K* group showed the opposite configuration. These oculomotor signatures index covert goal competition and align with the behavioral crossover.

### Phasic Pupil Dilation Tracks the Pre-Switch Control State

We next tested whether stimulus-locked phasic dilation in the block immediately preceding a choice differentiated upcoming switches from stays, and whether this depended on WM capacity. For each subject we computed ‘switch-minus-stay’ pupil trace after blink correction, then averaged within WM capacity groups. As shown in Figure 3C, low-*K* traces exhibited an early positive difference that rose shortly after stimulus onset and remained elevated with a significant cluster from 110-690 ms after trial onset, cluster-corrected *p* < .05. In contrast, high-*K* difference traces remained near zero throughout the trial. A direct between-group test on the difference traces was significant from 180-620 ms with low-*K* greater than high-*K*.

Pupil dynamics therefore link physiology to behavior and gaze control. Low-*K* participants approached voluntary switches with heightened early dilation consistent with conflict-driven effort under reactive control, rather than mere disengagement. High-*K* participants did not show this early elevation, which aligns with stronger goal-shielding and a tendency to initiate switches from a relatively well-focused state under proactive control.

### Impact of Decision Autonomy on Post-Selection Performance: Free versus Forced-Choice

To investigate how decision autonomy shapes the *consequences* of task selection, we compared commission rates on the subsequent *n*+1 block between the free-choice and forced-choice sessions, considering individual differences in WM capacity. This analysis directly addresses whether self-determined choice, as opposed to externally imposed task order, interacts with WM capacity in shaping goal maintenance, as predicted by theories of proactive and reactive control.

Across participants, the relation between WM and post-selection lapse rate diverged by decision autonomy. In the free-choice session (Figure 4A), higher WM was strongly associated with fewer commission errors on the *n*+1 blocks, *r*(27) = −.54, CI_95%_ [−.75, −.22], *p* < .001. In the forced-choice session (Figure 4B), the association was smaller and not significant, *r*(27) = −.27, CI_95%_ [−.58, .10], *p* = .150. Thus, when participants freely determined the next task to perform, individual differences in WM capacity predicted the quality of goal-shielding on the following block, whereas externally specifying the next task weakened this link, consistent with partial substitution of external goal specification for internal maintenance among lower WM capacity participants.

**Figure 4.**
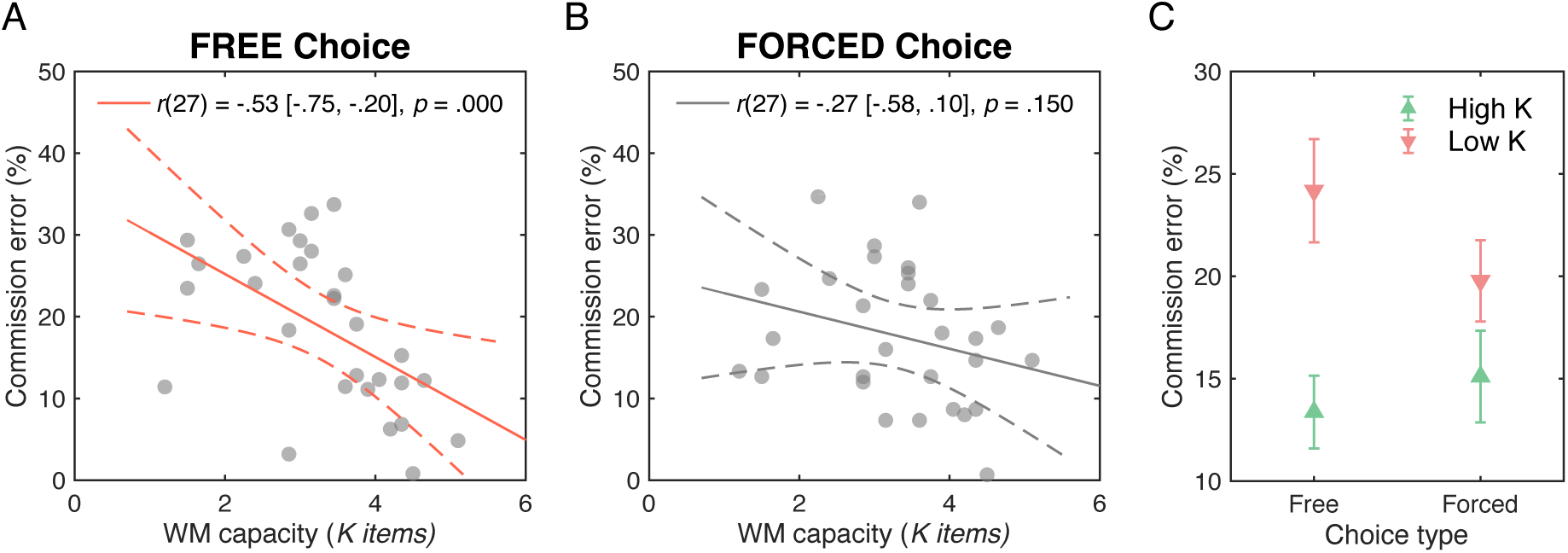
Decision autonomy selectively benefits lower working memory capacity individuals under forced-choice. Scatter plots of commission error rates on post-choice blocks versus individual working memory (WM) capacity estimates for (A) free-choice session and (B) forced-choice session. Solid and dashed lines represent best linear fit and 95% confidence intervals, respectively. (C) Median-split WM capacity group effect on post-choice block commission error rates by session type (free vs. forced-choice). Error bars represent standard error of the mean.

Median-split WM group means recapitulated these correlational results (Figure 4C). Low-*K* participants showed *lower* commission error rates under forced-choice, *M* = 20.1%, CI_95%_ [16.2, 24.0], than under free-choice, *M* = 24.4%, CI_95%_ [20.3, 28.6], whereas high-*K* participants were essentially comparable between forced-choice, *M* = 15.0%, CI_95%_ [10.6, 19.5], and free-choice sessions, *M* = 12.3%, CI_95%_ [8.8, 15.8]. This pattern yielded a significant two-way interaction effect between choice type (free vs. forced) and WM capacity group (high-*K* vs. low-*K*) on the post-choice commission error rates, *F*(1, 27) = 5.14, *p* = .032, η_p_² = .16, alongside a robust capacity group main effect, *F*(1, 27) = 7.99, *p* = .009, η_p_² = .25, and no significant main effect of choice type, *F*(1, 27) = 0.97, *p* = .333, η_p_² = .04. Externally imposing the upcoming task therefore selectively improved post-selection performance for low-*K* participants, consistent with the idea that enforced specification reduces goal competition and substitutes for weaker proactive shielding, while high-*K* participants who sustain stronger goal representations performed similarly regardless of autonomy.

We further tested whether autonomy influenced the classical task-switching cost and whether this effect was moderated by WM capacity (Figure 5). A three-way mixed ANOVA with factors of choice type (free vs. forced) × post-selection block (post-switch vs post-stay) × WM capacity group (high-*K* vs low-*K*) on commission errors revealed a reliable capacity group main effect, *F*(1, 27) = 8.09, *p* = .008, η_p_² = .23, as well as post-selection block main effect, *F*(1, 28) = 4.79, *p* = .037, η_p_² = .15, confirming a switch cost. Choice type did not show a significant main effect, *F*(1, 28) = 0.97, *p* = .333, η_p_² = .03. The three-way interaction was not significant, *F*(1, 27) = 0.05, *p* = .829, η_p_² = .00. Importantly, the choice type × WM capacity group interaction replicated the autonomy effect described above, *F*(1, 27) = 4.83, *p* = .037, η_p_² = .15, with lower lapse rates for low-*K* under forced-choice and comparable performance for high-*K* across sessions. The choice type × post-selection block interaction was not significant, *F*(1, 28) = 0.19, *p* = .662, η_p_² = .01, whereas the WM capacity group × post-selection block interaction was, *F*(1, 27) = 4.92, *p* = .035, η_p_² = .15, indicating that the switch cost was observed primarily in the low-*K* group.

**Figure 5.**
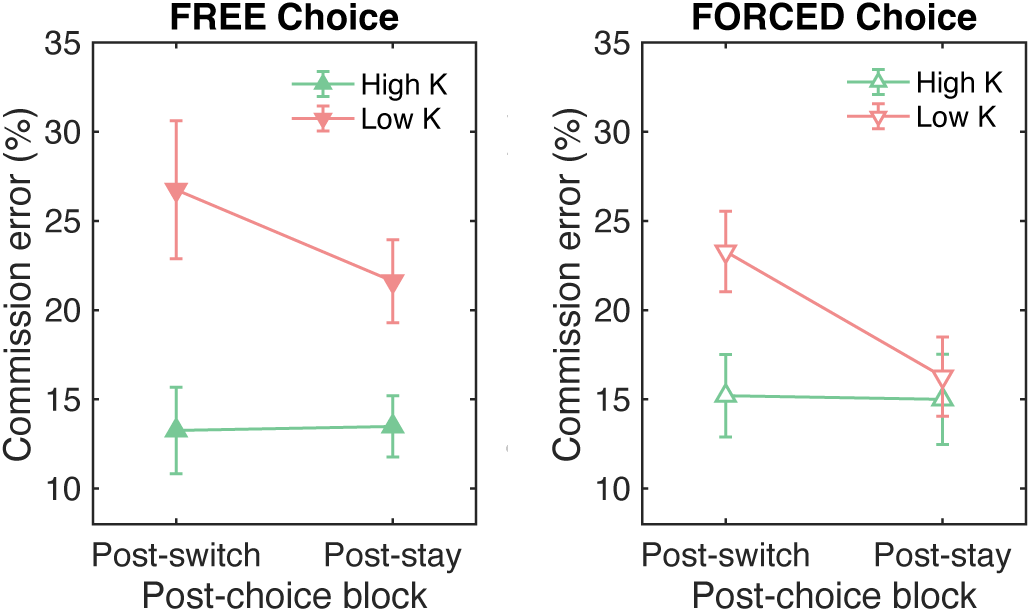
Post-selection commission error rates plotted separately for choice type (free vs. forced-choice), as a function of post-selection block (post-switch vs post-stay), and working memory capacity group (high *K* vs low *K*). Switch costs persist across decision contexts but overall lapse rates vary by autonomy and capacity. Error bars represent standard error of the mean.

Together, these findings show that decision autonomy does not change the size of the switch cost per se, but shifts the overall lapse level selectively for low-*K* participants. Forced selection lowered their lapse rates without eliminating the cost of switching. This pattern is consistent with the broader hypothesis that lower WM capacity is linked to reactive control under autonomy, whereas higher capacity supports proactive shielding that stabilizes post-selection performance across decision contexts.

## Discussion

The present study examined a goal-competition account of voluntary task switching by asking whether momentary lapses forecast the decision to abandon the current goal, and whether this prospective link depends on WM capacity. Across behavioral, oculomotor, and pupillometric indices, voluntary switches were preceded by capacity-dependent differences in the control state of the ongoing block. Individuals with lower WM capacity were more likely to switch after lapse-prone blocks, whereas those with higher capacity tended to switch from relatively well-focused states. This qualitative dissociation is difficult to reconcile with accounts that treat lapses as uniform control failures. Instead, it points to lapses as a window into competition among concurrently active goals, with outcomes contingent on individual control resources (Robertson et al., 1997; Smallwood & Schooler, 2015).

The free-choice analyses provide the central evidence for this pattern. Lapse-related measures in block *n* prospectively predicted whether participants voluntarily switched on block *n+1*, and this predictive relation depended on WM capacity. Specifically, commission errors, defined as failures to withhold responses on no-go trials, indexed response-level lapses, whereas swap errors, defined as responses to the currently irrelevant category, provided a more direct marker of task-level goal confusion. In the simultaneous GLME, both *Commission × K* and *Swap × K* interactions were reliable, indicating that the relation between prior-block errors and subsequent switching became increasingly negative as WM capacity increased. Converging evidence from eye movements and pupillometry supported this interpretation. Lower-capacity participants showed greater distractor-directed oculomotor bias and heightened early pupil dilation prior to voluntary switches, whereas higher-capacity participants exhibited a more target-aligned pre-switch state. Together, these findings indicate that the same overt act of switching can arise from qualitatively different internal control states, aligning a goal-competition perspective with contemporary theories of cognitive control that emphasize individual differences in the balance between stability and flexibility (Braver, 2012; Kiesel et al., 2010; Monsell, 2003). More broadly, this pattern also speaks to conflict adaptation in multitasking contexts, by showing that the control state preceding a task choice can contain separable signatures of response-level and task-level conflict. In the present task, commission errors indexed failures to withhold a response on no-go trials, whereas swap errors indexed responses guided by the currently irrelevant task category. Thus, commission errors reflected a more general lapse in response control, whereas swap errors provided a more specific marker of competition between simultaneously active task goals. The fact that both measures prospectively predicted voluntary switching in a WM-dependent manner suggests that task choice is shaped not only by the occurrence of performance failures, but also by how individuals manage competition between the currently relevant task set and an alternative task set. From this perspective, stability and flexibility may not constitute a fixed trade-off across individuals.

Higher WM capacity may allow participants to maintain a stable task representation while still flexibly initiating a voluntary switch, whereas lower WM capacity may make switching more likely to emerge as a reactive response to instability and goal competition.

A critical question is whether this capacity-dependent pattern reflects differences in underlying control dynamics or distinct strategic responses to performance failures. A purely strategy-based account might treat lapses as a generic failure signal that increases the subjective appeal of switching. Under this view, capacity difference would arise from different decision thresholds, such that low-capacity individuals abandon tasks more readily when performance deteriorates, whereas high-capacity individuals persist longer in the hope of recovery. The present data suggest a more specific interpretation. Commission and swap errors captured separable aspects of the preceding control state, and each predicted subsequent task choice in interaction with WM capacity. Thus, the critical effect was not simply that poor performance increased switching, but that the prospective meaning of error-related signals depended on the individual’s WM capacity. Moreover, the oculomotor and pupillary findings indicate that switches in lower-capacity participants were preceded by heightened competition from the currently irrelevant task set, rather than by behavioral failure alone. In contrast, higher-capacity participants tended to switch from stable, target-focused states and were less likely to switch after error-prone blocks. This suggests that WM capacity shapes whether voluntary goal shifts emerge from reactive control under conditions of competition or from more proactive exploration initiated from a stable control state.

These findings align with proposals that lapses reflect episodes when alternative goals gain traction against the prioritized task set. In the present task, every display contained both a relevant and an irrelevant category, creating a setting in which response-level failures (i.e., commission errors) and task-level goal confusion (i.e., swap errors) could be dissociated. The fact that these measures were related but nonredundant, and that both showed capacity-dependent prospective relations with switching, supports the broader claim that lapses are not merely passive losses of control. Rather, they can reveal transient shifts in the balance between competing task representations. The early pupil dilation observed in lower-capacity participants further suggests conflict-driven effort under heightened competition rather than disengagement, consistent with accounts linking pupil responses to noradrenergic arousal and adaptive control (Aston-Jones & Cohen, 2005; Joshi & Gold, 2020; van der Wel & van Steenbergen, 2018). From this perspective, voluntary switching reflects not only a decision to change tasks, but also the control state from which that decision emerges. This may help explain why the WM-dependent pattern was clearer for switch decisions than for stay decisions. A switch provides a relatively discrete marker of abandoning the current task set, whereas a stay decision may reflect multiple latent states, such as stable maintenance, reactive re-engagement after instability, or constraints from the remaining task schedule. Thus, the present findings do not imply that WM capacity is irrelevant for staying, but rather that voluntary switches may provide a cleaner behavioral readout of whether or how goal competition leads to reactive escape from the current task set or to deliberate flexible updating from a stable state.

The dual-mechanisms of control (DMC) framework provides a principled interpretation of this capacity-dependent crossover. Higher-capacity individuals appear to rely more on proactive control, allowing them to maintain task-relevant goal representations and initiate switches from relatively stable states. Lower-capacity individuals, in contrast, rely more on reactive control, such that competing task representations more readily influence behavior before a switch occurs. This extends DMC accounts to voluntary behavior by suggesting that task choice is shaped not only by external task demands, but also by the control state from which the choice emerges (Braver, 2012; Kane & Engle, 2003; Redick, 2014; Gonthier et al., 2016). Lower-capacity individuals should not be understood as simply having weaker motivation to persist. Rather, their performance suggests a reduced ability to shield the current task set from competing alternatives. As a result, task-level competition may become more likely to appear as swap errors, distractor-directed gaze, and heightened pupil-linked effort before switching. In contrast, higher-capacity individuals appear better able to maintain stable goal representations in the presence of competing alternatives, allowing voluntary switches to reflect more deliberate exploration from a well-controlled state rather than reactive escape from goal competition. Neuroimaging findings linking proactive control to sustained prefrontal activity during goal maintenance further support this interpretation (Braver et al., 2009).

The autonomy manipulation further clarifies this distinction. Externally imposed task selection selectively reduced post-selection lapse rates for lower-capacity participants without eliminating switch costs, suggesting that external goal specification can scaffold control when internal maintenance is limited. In contrast, higher-capacity participants performed consistently regardless of autonomy, indicating that robust internal goal representations reduced the need for external support. This pattern suggests that the reactive control profile observed in lower-capacity individuals reflects a context-sensitive limitation in goal maintenance rather than a fixed tendency to perform poorly or abandon tasks. When the next task was specified externally, lower-capacity participants no longer needed to maintain and evaluate competing task options to the same extent, and their lapse rates improved. This is consistent with work on cognitive offloading and situational support, which shows that external structure can stabilize performance for individuals with weaker internal control while having minimal impact on those with stronger control (Duckworth et al., 2016; Inzlicht et al., 2014; Risko & Gilbert, 2016).

Methodologically, the present study leveraged a prospective approach, relating upcoming task choices to the control state in the preceding block rather than inferring control solely from post-switch costs. The convergence of behavior, gaze, and pupil measures traces a coherent path from internal competition to overt task choice. Framed within exploration-exploitation theory, our results imply that criteria for leaving a task depend jointly on the current control state and the individual’s WM capacity to maintain task goals (Daw et al., 2006; Schulz & Gershman, 2019; Wilson et al., 2014). This perspective aligns with foraging accounts in which agents arbitrate between persisting with a known option and sampling alternatives, a computation supported by frontal valuation and control circuits (Kolling et al., 2012). Within this framework, lower-capacity individuals may engage exploration through reactive escape from mounting goal competition, whereas higher-capacity individuals may engage exploration through more deliberate sampling from stable goal states.

Several limitations point to directions for future research. First, although the present findings revealed a consistent capacity-dependent pattern across behavioral, oculomotor, and pupillometric measures, the sample size was only modest for estimating individual-difference effects and cross-level interactions. The observed capacity-dependent crossover should therefore be interpreted as initial evidence for a theoretically informative pattern rather than as a precise estimate of the population-level form or magnitude of the effect. Future studies with larger samples will be important for evaluating the robustness, generalizability, and limits of this pattern. Second, the forced-choice session used a single fixed sequence for all participants.

Consequently, differences between free- and forced-choice sessions may partly reflect incidental sequence statistics, such as switch rate or run-length distribution, in addition to the locus of control. Future studies could better align sequence properties across conditions by constructing forced-choice sequences from free-choice patterns produced by other participants.

Future work could also use more temporally resolved measures to track the emergence of competing task representations. Direct neural measures, such as the N2pc component measured from electroencephalogram (EEG), could test whether distractor-directed selection increases before voluntary switches (Luck & Hillyard, 1994; Feldmann-Wüstefeld & Vogel, 2019), and multivariate <u>neural decoding</u> analyses could test whether the irrelevant task set becomes decodable during the pre-switch interval (Waskom et al., 2014). Complementarily, mouse-tracking could provide a continuous behavioral readout of how conflict in control or decisional states unfolds before it is resolved in a final response. Prior work has shown that mouse trajectories can reveal covert competition among internal representations as decisions evolve over time (Park, 2025; Park & Zhang, 2026, 2022). Applied to voluntary task switching, trajectory-based measures could clarify how competing task goals are expressed in action across spatial bias, movement timing, and decisional consistency as participants approach a switch or stay decision.

In conclusion, voluntary task switches may be preceded by distinct internal states reflecting the management of goal competition. By linking prospective lapse-related behavior, oculomotor dynamics, and pupil-linked effort to WM capacity, these findings suggest that lapses are not homogeneous control failures, but informative markers of an individual’s control mode. The capacity-dependent pattern observed here highlights the importance of individual differences for theories of flexible, goal-directed behavior and suggests that WM capacity shapes how the system resolves competition between stability and flexibility. This perspective resonates with broader theoretical developments linking mind wandering to exploration-exploitation trade-offs, wherein off-task states may reflect the control system’s attempt to optimize resource allocation across competing goals (Hills & Hertwig, 2010). Understanding how distinct control modes shape the route to switching has practical relevance for designing environments and interventions that calibrate autonomy and structure to cognitive resources, supporting sustained performance while acknowledging diversity in cognitive control profiles.

## Acknowledgement

This research was supported by Office of Naval Research grant N00014-22–1-2123 to E.K.V. and Multidisciplinary University Research Initiatives (MURI) grant N00014-23-1-2768 to E.K.V. and M.D.R.

## Notes

### Competing Interest Statement

The authors have declared no competing interest.

### Summary of Updates

The paper has been revised to clarify the continuous performance task framework and to define commission errors and swap errors more explicitly. The Introduction now provides a clearer rationale for linking transient control failures during sustained task performance to later voluntary task choices. The statistical models were revised to include the relevant random effects, and additional checks were added to examine the overlap between error types and the robustness of the findings after excluding participants with highly patterned switching behavior. The Discussion was updated to sharpen the interpretation of distinct control states, working memory capacity, and voluntary control. A Transparency and Openness statement was added. Citations and references were corrected.

